# Machine learning enables detection of early-stage colorectal cancer by whole-genome sequencing of plasma cell-free DNA

**DOI:** 10.1101/478065

**Authors:** Nathan Wan, David Weinberg, Tzu-Yu Liu, Katherine Niehaus, Daniel Delubac, Ajay Kannan, Brandon White, Eric A. Ariazi, Mitch Bailey, Marvin Bertin, Nathan Boley, Derek Bowen, James Cregg, Adam M. Drake, Riley Ennis, Signe Fransen, Erik Gafni, Loren Hansen, Yaping Liu, Gabriel L Otte, Jennifer Pecson, Brandon Rice, Gabriel E. Sanderson, Aarushi Sharma, John St. John, Catherina Tang, Abraham Tzou, Leilani Young, Girish Putcha, Imran S. Haque

**Author notes:** Co-first Author (Nathan Wan, David Weinberg). Co-senior Author (Girish Putcha, Imran S. Haque).

## Abstract

**Background:** Blood-based methods using cell-free DNA (cfDNA) are under development as an alternative to existing screening tests. However, early-stage detection of cancer using tumor-derived cfDNA has proven challenging because of the small proportion of cfDNA derived from tumor tissue in early-stage disease. A machine learning approach to discover signatures in cfDNA, potentially reflective of both tumor and non-tumor contributions, may represent a promising direction for the early detection of cancer.

**Methods:** Whole-genome sequencing was performed on cfDNA extracted from plasma samples (N=546 colorectal cancer and 271 non-cancer controls). Reads aligning to protein-coding gene bodies were extracted, and read counts were normalized. cfDNA tumor fraction was estimated using IchorCNA. Machine learning models were trained using k-fold cross-validation and confounder-based cross-validation to assess generalization performance.

**Results:** In a colorectal cancer cohort heavily weighted towards early-stage cancer (80% stage I/II), we achieved a mean AUC of 0.92 (95% CI 0.91-0.93) with a mean sensitivity of 85% (95% CI 83-86%) at 85% specificity. Sensitivity generally increased with tumor stage and increasing tumor fraction. Stratification by age, sequencing batch, and institution demonstrated the impact of these confounders and provided a more accurate assessment of generalization performance.

**Conclusions:** A machine learning approach using cfDNA achieved high sensitivity and specificity in a large, predominantly early-stage, colorectal cancer cohort. The possibility of systematic technical and institution-specific biases warrants similar confounder analyses in other studies. Prospective validation of this machine learning method and evaluation of a multi-analyte approach are underway.

## Introduction

Despite the public health emphasis on population-level cancer screening in recent decades, adherence remains lower than desired [1], and cancer is often detected too late for successful treatment. For example, nearly 60% of colorectal cancer (CRC) cases, and approximately 80% of pancreatic cancer cases, are detected after regional or distant metastases [2]. Current cancer screening methods are often invasive, inconvenient, expensive, and/or have suboptimal clinical performance (i.e., sensitivity or specificity), particularly for early-stage disease and precancerous lesions [3].

Recently, blood-based screening tests for cancer have been proposed in an effort to address some of the aforementioned challenges. One key area of both academic and commercial interest is circulating cell-free DNA (cfDNA), which includes both tumor-derived DNA (so-called “circulating tumor DNA”, or ctDNA) and DNA derived from non-tumor cells, such as hematopoietic and stromal cells, to supplement or replace existing cancer screening methods.

Different screening approaches using cfDNA are being explored, and some have hypothesized that ctDNA-only based “liquid biopsies” may enable sensitive and specific early detection of cancer ([4–7]. ctDNA has unique characteristics of tumor DNA, such as cancer-associated mutations, translocations, and/or large chromosomal copy number variants (CNVs), not typically present in the cfDNA of healthy patients [8]. In addition, ctDNA fragments appear to be shorter on average than cfDNA found in healthy subjects [9]. However, others have questioned whether such an approach is feasible for routine screening, given biological (e.g., clonal hematopoiesis of indeterminate potential (CHIP)), technical (e.g., limits of detection and variable levels of tumor fraction (TF) observed in cancer patients), and practical (e.g., blood volume requirements and cost) considerations [10–12]. In patients with cancer, ctDNA generally represents a small fraction of all cfDNA, ranging from ≥5-10% in late-stage disease to ≤0.01-1.0% in early-stage disease, and even lower in premalignant conditions [13]. These limitations are particularly important in early-stage cancer when the tumor is small and the shedding of DNA into the blood may be minimal. Indeed, many previous cfDNA studies have had stage distributions meaningfully different from those seen in screening populations [14–16].

An alternative to detection based solely on ctDNA is to look more broadly at cfDNA—both tumor derived and non-tumor derived—and changes that early-stage cancer may induce in blood. There is growing evidence of interactions between cancerous cells and other cells, including fibroblasts, platelets, and immune cells, especially within the tumor microenvironment. These include findings of “tumor education”, such as changes in gene expression that may reflect interaction with a tumor and/or ingestion of tumor-related molecules [17]. For example, platelets in patients with cancer harbor different patterns of messenger RNA (mRNA) than platelets in healthy individuals [18]. There are also reports of changes in immune-cell apoptosis patterns in patients with cancer [19], suggesting global changes in hematopoietic cell populations that may reflect altered physiological states. For instance, low relative levels of circulating lymphocytes versus monocytes may be correlated with poor cancer prognosis [20]. It is possible to detect such changes in cell populations from cfDNA because cfDNA fragmentation and methylation patterns can recapitulate expected cellular epigenetic states [15,16,21–23].

Because it is still unknown to what extent circulating cells in patients with early-stage cancer are educated by the tumor microenvironment (i.e., how changes in cellular state are explicitly reflected in the billions of base pairs of cfDNA), the ability to identify disease-relevant patterns in cfDNA requires unbiased methods that can identify patterns in high-dimensional space. Given a large enough sample size, machine learning (ML) may provide a toolset by which to learn disease-related patterns from whole-genome signals directly from patients with and without early-stage cancer. However, the primary challenge in an assay with many measured variables is to identify relevant, low-dimensional features that generalize to the screening population [24,25]. As a corollary, it is necessary to mitigate potential confounding variables, defined as variables that are correlated with the clinical label which in this case is the disease label. For example, batch effects or institutional processing effects can be a significant variable that correlates with non-cancerous and cancerous samples.

Here we develop and implement a computational approach for representing and learning associations between cfDNA profiles and cancer status, with a focus on the importance of accounting for known confounding variables. Using this approach, we report classification results for a large cohort of non-cancer controls and early-stage CRC patients.

## Materials and Methods

### Sample collection

Human EDTA plasma samples were acquired from 546 patients diagnosed with CRC (Table 1). As controls, plasma samples from 271 unique patients without a current CRC diagnosis were also acquired. In total, 817 de-identified plasma samples were collected from institutions and commercial biobanks from Europe and the United States. Patient age, gender, and cancer stage (where available) were obtained for each sample. Samples were included in the intended use (IU) age range analysis for CRC only if the patient’s age at time of collection was known to be between 50 and 84, inclusive. Plasma was stored at –80°C.

**Table 1:**
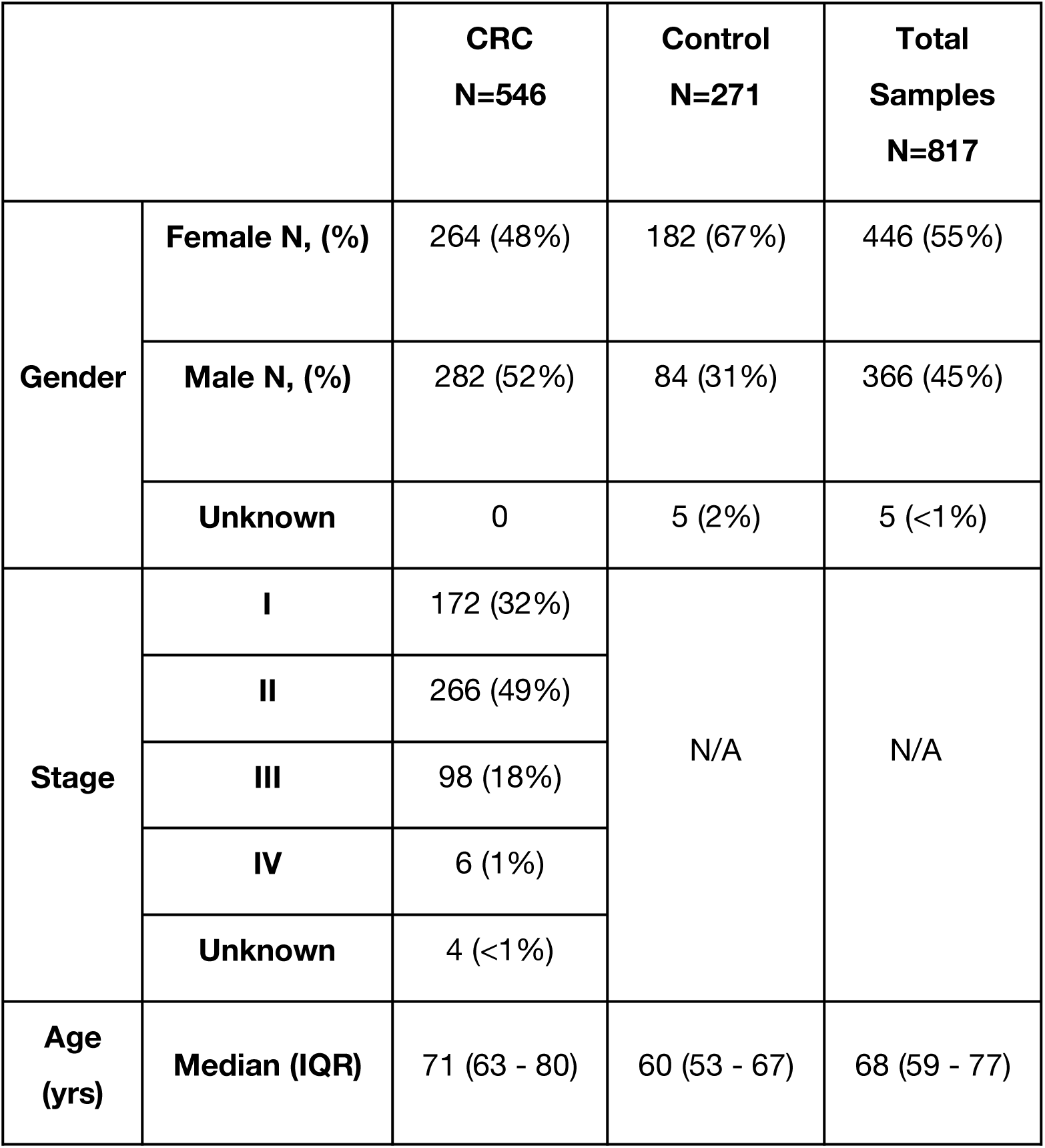
Clinical characteristic and demographics of CRC patients and non-cancer controls

### Laboratory processing, bioinformatics, and featurization

Detailed descriptions of laboratory processing and sequencing, bioinformatics analysis, data preprocessing, classifier training, and validation methods (including measuring and controlling for confounding factors) are provided in the Supplemental Methods.

Briefly, cfDNA was extracted from 250 µl plasma using the MagMAX cfDNA Isolation Kit (Applied Biosystems), converted into libraries using the NEBNext Ultra II DNA Library Prep Kit (New England Biolabs), and paired-end sequenced on the Illumina platform. Reads were aligned to the human genome using BWA-MEM 0.7.15 [26]; all datasets passing quality control (based on acceptable GC bias, sufficient number of reads, and no evidence of contamination or sample swaps) continued to featurization. Aligned reads were transformed into per-sample feature vectors by counting the number of fragments appearing in protein-coding genes. Features were normalized per-sample by dividing by the trimmed mean (excluding top and bottom 10% of counts) over all features and applying Loess GC bias correction [27]. Categorical features used in certain experiments (binned age, sex, and institution) were featurized using a one-hot encoding. TF was estimated in each sample using IchorCNA [28] from read counts in 1-Megabase (Mb) bins across the entire genome.

### Model training

ML models were trained and evaluated using cross-validation (CV) procedures as follows. Each feature, which is a preprocessed read count, was standardized by subtracting the mean and dividing by the standard deviation after large outliers were replaced with 99th percentile value. Dimension-reduction methods including principal component analysis (PCA) and truncated singular-value decomposition (SVD) were then optionally applied to the standardized data. Two classification methods were considered for training (logistic regression and support vector machine (SVM)) with hyperparameters chosen based on random search. The best model was selected based on k-fold CV, and the methods were subsequently applied to other CV procedures. All methods were implemented by Scikit-learn [29].

### Validation and confounder control

Five different CV schemes were used to obtain estimates of model performance. All CV procedures shared in common the partitioning of the data into multiple independent subsets, or “folds,” with individual folds held out and used to assess the performance of models trained on the remaining data. The principal difference among CV procedures was how individual samples were partitioned into folds. The procedures included k-fold, in which samples were partitioned at random (stratified by class label of cancer or not cancer); binned-age, in which partitions were defined based on age; k-batch, in which partitions were defined by processing batch; balanced k-batch, in which partitions were defined by processing batch with additional downsampling to stratify by institutional source; and ordered k-batch, in which samples were partitioned by date of laboratory processing. Further explanation of each method is provided in the Supplemental Methods.

Five folds (k=5) were used for CV of all models except for binned-age (which has a fixed number of bins). Reported performance metrics are mean area under the receiver operating characteristic curve (AUC) and mean sensitivity at 85% specificity, each calculated across all test folds.

## Results

Paired-end whole-genome sequencing (WGS) was performed on plasma cfDNA obtained from 271 non-cancer control subjects and 546 CRC patients (Table 1). The patient population was approximately equally split by gender (55% female, 45% male), and consisted of 80% early-stage (stages I and II) patients. The non-cancer control population skewed younger (median age = 60; interquartile range [IQR] = 53-67) than the cancer population (median age = 71; IQR = 63-80, p<0.01, Mann-Whitney U-test) (Table 1B).

WGS data were converted into input features for the classification model by counting the number of fragments overlapping each annotated protein-coding gene (i.e., each gene corresponded to a single bin) and then normalizing to account for feature length, mappability, read depth, and sequence-content biases. The gene-based featurization was designed to simultaneously capture both copy number changes as well as epigenetic signals reflected in cfDNA fragmentation patterns across genes [15].

Before assessing classification performance, models were trained using confounding variables as inputs to validate our CV stratification methods. In k-fold CV, binned age, batch, processing date, and institution confounders achieved mean AUCs of 0.71, 0.72, 0.69, and 0.87, respectively, when tested individually as the only input features to the classification model (Table 2, Supplemental Figure 1). When evaluated using CV methods tailored specifically to address them, these same input features (i.e., confounder variables) gave at-chance predictive performance (i.e., mean AUC = 0.50), demonstrating that binned-age, k-batch, balanced-k-batch, and ordered k-batch CV effectively control for their respective confounder variables (Table 2, Supplemental Figure 1).

**Table 2:**
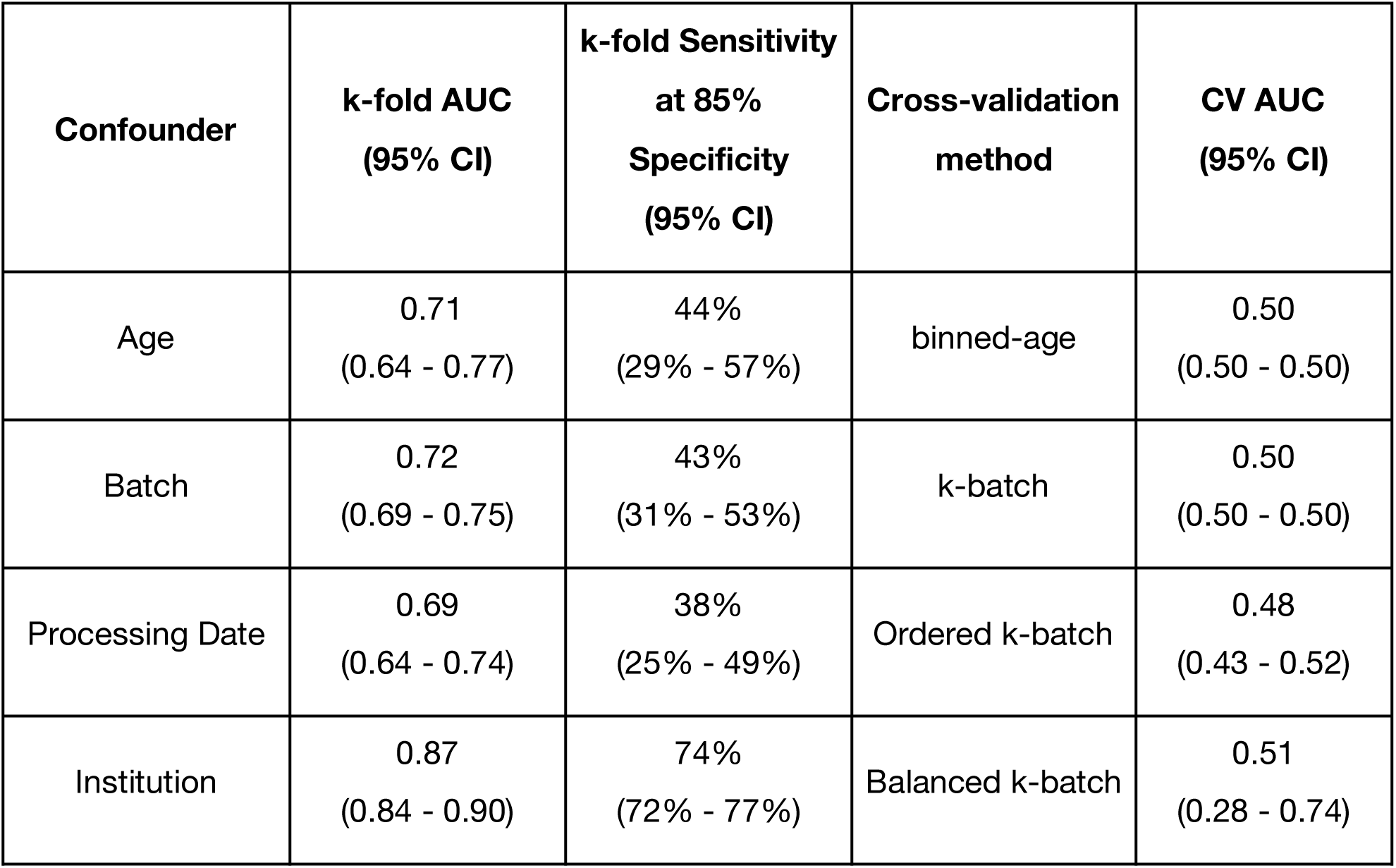
Performance evaluation of known confounders alone to predict cancer with either k-fold or the CV procedure designed to control for the confounder. Confidence intervals are calculated from bootstrapped distributions of the metric across folds.

**Figure 1A/B:**
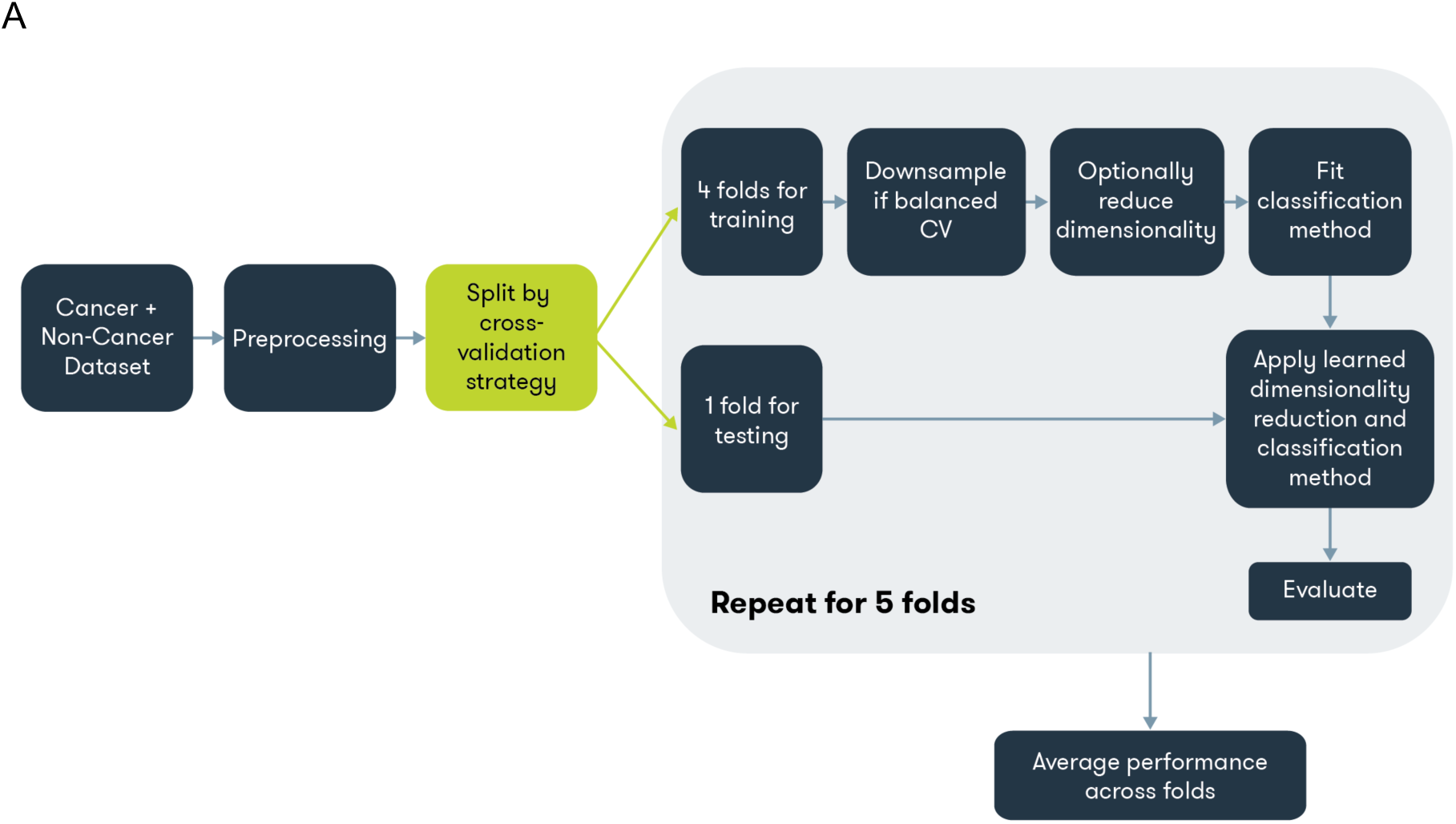
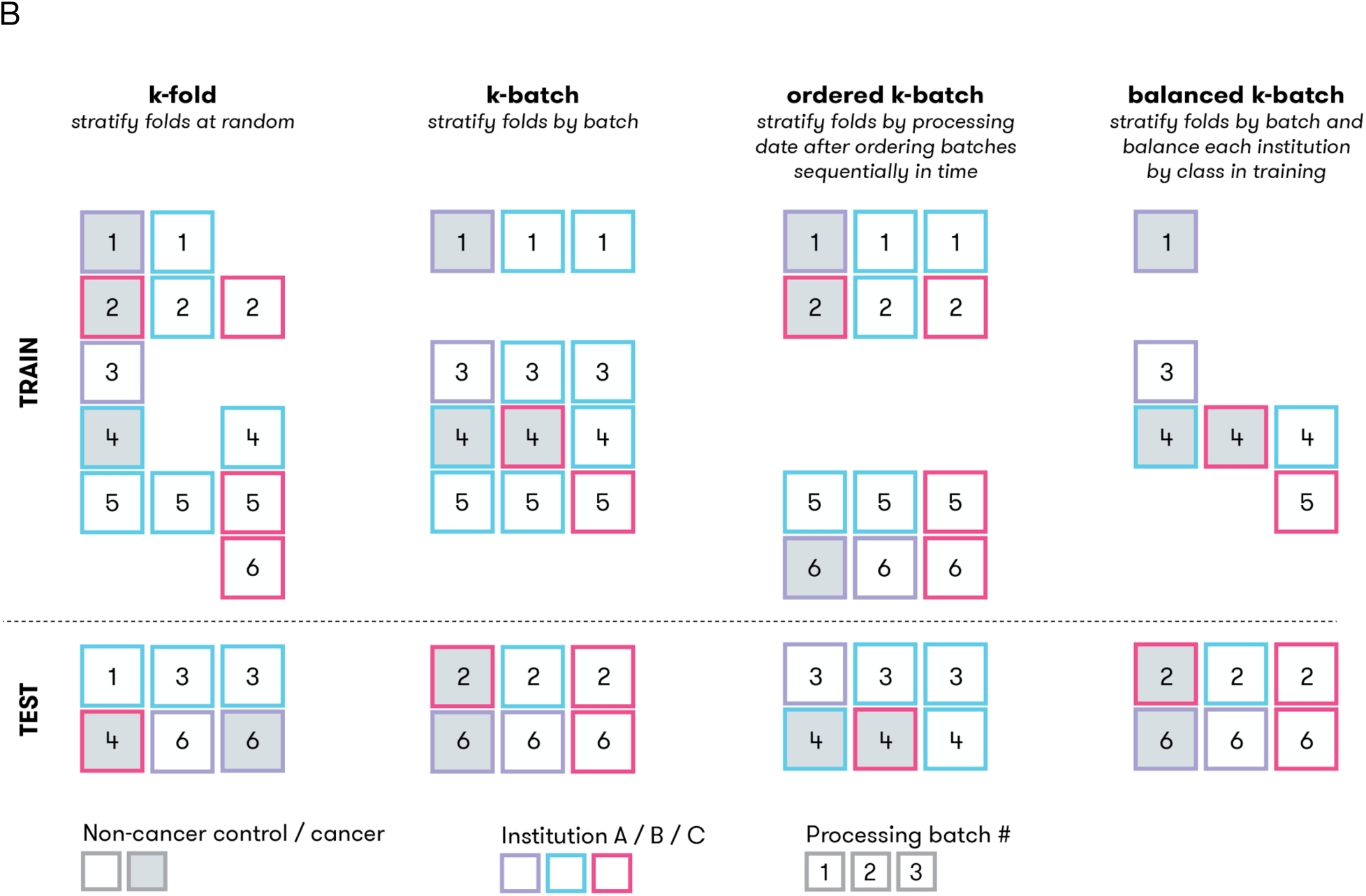
Model training overview and CV procedures. **A)** All methods were trained on k-fold, and the best performing method was chosen to train models for the other cross-validation procedures. Diagram describes individual steps in common to all methods. Models are trained on a given dataset and set of methods (i.e., dimension reduction and classification) and then evaluated, resulting in a performance estimate. **B)** Illustration of CV procedures for k-fold, k-batch, ordered k-batch, and balanced k-batch. Each square represents a single sample, with the fill color indicating class label, the border color representing a confounding factor like institution, and the number indicating processing batch. Each column represents a possible fold constructed for the given CV procedure. The dashed line separates the test set of samples held out from the training set.

After initial model selection via k-fold CV performance, we additionally applied each previously introduced CV procedure to the same methods to estimate the generalizability of performance when controlling for particular confounder variables individually (Table 3). The method selected by k-fold CV performance was no dimensionality reduction and SVM classification. Evaluation by standard k-fold CV achieved a mean AUC of 0.92 (95% bootstrap confidence interval (CI) of 0.91-0.93) with a mean sensitivity of 85% (95% CI = 83-86%) at 85% nominal specificity. Using binned-age CV to control for age achieved mean AUC of 0.91 (95% CI = 0.89 - 0.94) with a mean sensitivity of 79% (95% CI = 73-87%) at 85% specificity. We controlled batch-to-batch technical variability using k-batch CV and process variability using ordered k-batch CV, which achieved mean AUC of 0.91 (95% CI = 0.88 - 0.94) and 0.90 (95% CI = 0.83 - 0.94) and sensitivity at 85% specificity of 85% (95% CI = 80 - 89%) and 73% (95% CI = 53 - 88%), respectively. The larger variance observed in ordered k-batch may be attributed (at least in part) to higher standard deviation in test fold sizes (80.8) when compared to standard deviation of test folds of k-batch (35.0) (Supplementary Table 1). Finally, we applied balanced k-batch CV to control for possible institution-specific differences in population or sample handling. Despite training on a significantly reduced dataset (average of 263.6 samples per fold in training versus 653.6 samples per fold with k-fold or k-batch as seen in Supplementary Table 1), the balanced k-batch CRC model achieved a mean AUC of 0.83 (95% CI = 0.79 - 0.86) with a mean sensitivity of 71% (95% CI = 63-76%) at 85% specificity (Table 3). Figure 2 shows ROC curves for each CV procedure.

**Table 3:** CRC performance by cross-validation procedure in 50-84 year-old patients. Confidence intervals are calculated from bootstrapped distributions of the metric across folds.

| **Validation** | **Mean AUC**<br>**(95% CI)** | **Mean Sensitivity at 85% Specificity**<br>**(95% CI)** |
| --- | --- | --- |
| **k-fold** | 0.92<br>(0.91-0.93) | 85%<br>(83-86%) |
| **Binned-age** | 0.91<br>(0.89-0.94) | 79%<br>(73-87%) |
| **k-batch** | 0.91<br>(0.88-0.94) | 85%<br>(80-89%) |
| **Ordered k-batch** | 0.90<br>(0.83-0.94) | 73%<br>(53-88%) |
| **Balanced k-batch** | 0.83<br>(0.79-0.86) | 71%<br>(63-76%) |

**Figure 2:**
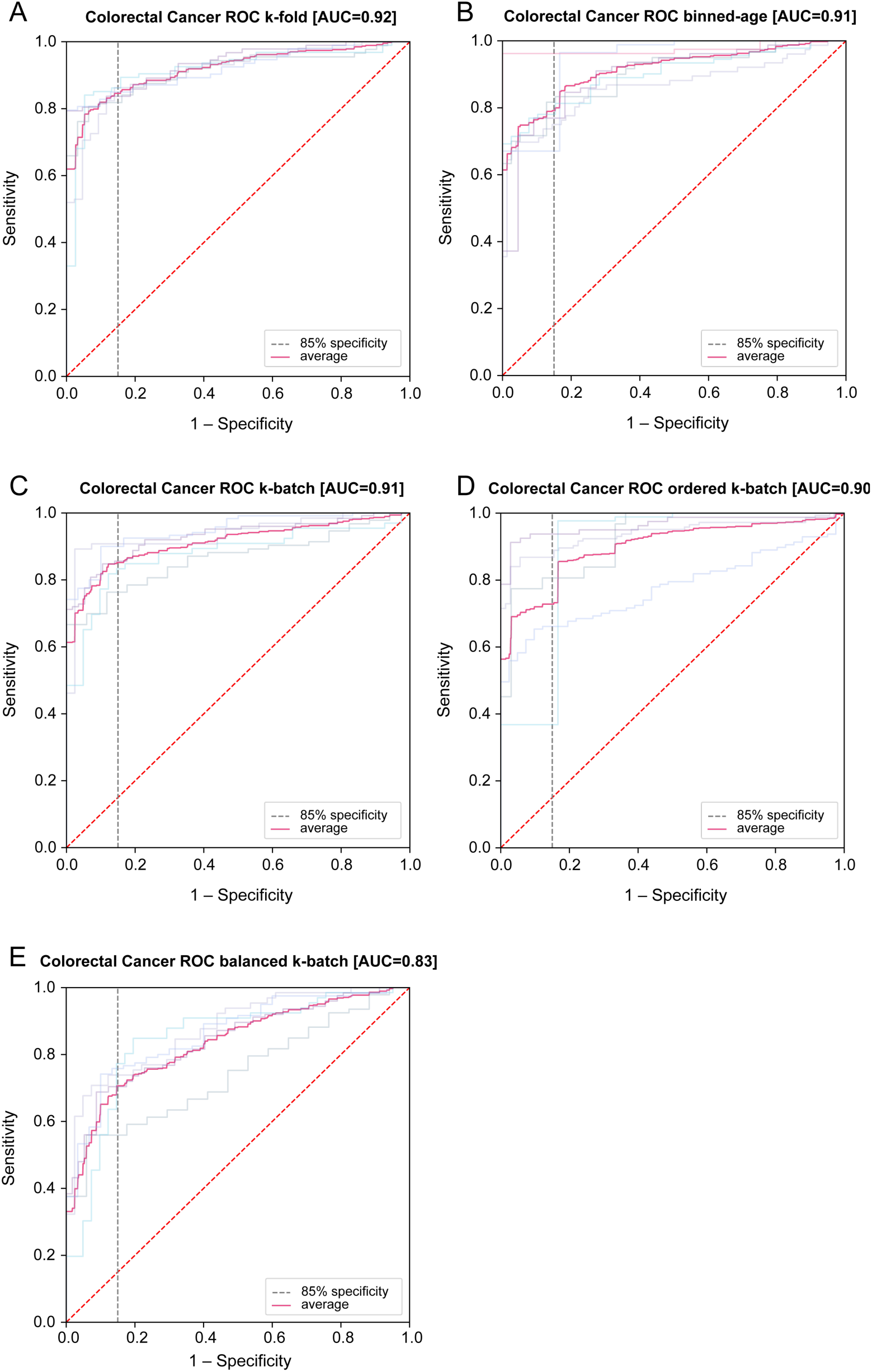
Colorectal cancer classification performance (ROC curves) by each cross-validation method. Average of all folds drawn in solid blue; random chance is represented as dashed red; ROCs for each fold drawn behind. A) k-fold, B) binned age, C) k-batch, D) ordered k-batch, and E) balanced k-batch

Additionally, we investigated the sensitivity of our method, trained using each CV procedure, to relevant clinical parameters. Figure 3A illustrates sensitivity as a function of clinical stage. All validation methods achieved similar distributions of sensitivity across stages I through III, and consistently classified stage IV cancer correctly. Stage II samples, which represent the majority of our data, performed consistently well. We also evaluated age, which is a known confounder. The AUC performance increased with age in nearly all validations (Figure 3B). Taken as a whole, the results are consistent with the general notion that cancer is an age-related disease. Performance for males and females was comparable across validation types (Figure 3C), even in spite of the observed imbalance in non-cancer controls (Table 1).

**Figure 3:**
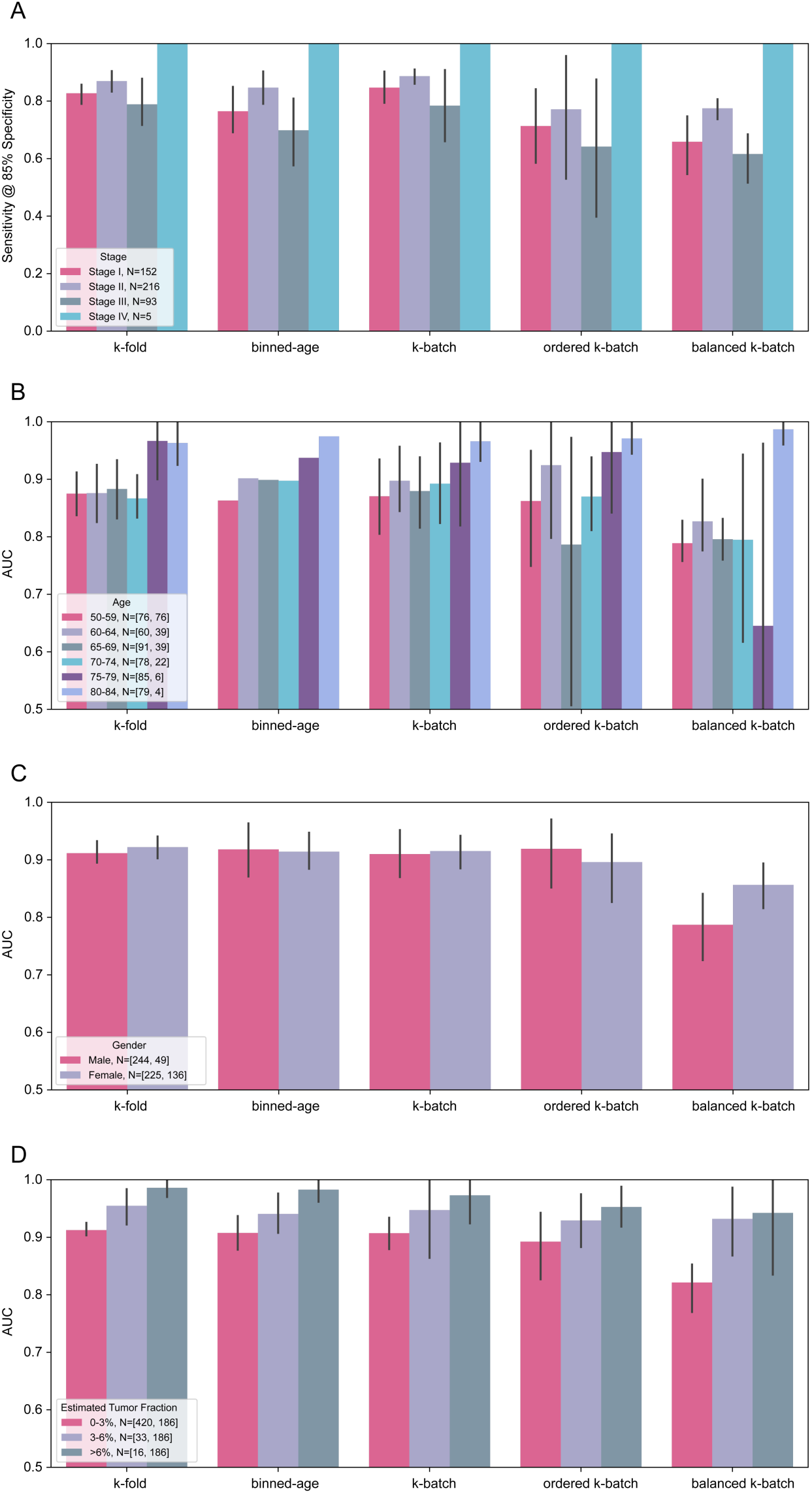
Classification performance for colorectal cancer within the IU age range across all validation methods. N is number of samples, [cancer, controls]. **A.** Sensitivity at 85% nominal specificity by CRC stage across all CV procedures **B.** AUC by age bins across all CV procedures **C.** AUC by gender across all CV procedures **D.** AUC by IchorCNA-estimated TF across all CV procedures

Tumor fraction (TF), defined as the fraction of cfDNA originating from tumor cells, has been implicated as a critical parameter for the design of blood-based cancer screens [4,6,12,30,31]. As high-depth mutation detection information is not available in our data, we estimated TF from observed copy number variation using IchorCNA [28]. The majority of control samples (98%) were estimated to have a TF below 3%, which is consistent with IchorCNA’s estimated limit of detection (Supplementary Figure 2).

Figure 3D displays our CRC model’s AUC as a function of IchorCNA-estimated TF. Observed performance declined with decreasing TF, which is consistent with the hypothesis that an ML-based method may be able to detect tumor-derived signal; however, performance remained better than chance even in the lowest TF bin. To investigate whether the ML model may detect signal beyond tumor-derived CNVs, we used IchorCNA-estimated TF alone to predict cancer. This method achieved AUC of 0.62 in the IU age range, lower than results from the ML model under any analyzed CV scheme (Table 3), consistent with the possibility that the ML model used non-tumor-derived signal (i.e., beyond IchorCNA-detectable CNVs) (Supplementary Figure 3).

To address decreased classifier performance due to smaller sample sizes in training (i.e., balanced k-batch), the CRC dataset was downsampled. Supplemental Figure 4 illustrates the non-linear relationship between the total number of samples used for training and the measured sensitivity. These results suggest that the lower performance observed using balanced k-batch is explained, at least in part, by the smaller size of the training dataset.

## Discussion

Our results show promising preliminary performance for early-stage (i.e., stages I and II) CRC detection using blood. To our knowledge, this multicenter, international study represents the largest study to date using only cfDNA WGS in patients for the early detection of CRC. We have demonstrated that it is possible to take an ML-based approach to learn the relationship between a patient’s cfDNA profile and cancer diagnosis, with 85% sensitivity at 85% specificity in CRC using standard k-fold cross-validation; application of other rigorous and novel CV strategies specifically designed to control for known confounding variables yielded 71-85% sensitivity at 85% specificity.

In this work, we focused our approach on cfDNA count profiles across the whole human genome (~3200 Mb) at relatively low depth (~9X), as opposed to existing liquid biopsy approaches that assess small regions (<2 Mb) of the genome at very high depth (~60,000X) to detect tumor-derived mutations. In particular, we applied ML methods to perform unbiased discovery of signals of varying origin that may inform on the presence of a tumor (including both tumor-derived CNVs as well as potentially non-tumor-derived signals such as changes in the epigenetic states of circulating immune cells) vis-à-vis focusing on only tumor-derived mutations. This parallels previous research in non-invasive prenatal testing (NIPT): Kim et al. demonstrated that an ML-based regression algorithm operating on genome-wide count data was able to accurately estimate fetal fraction in the cfDNA of pregnant women, without the detection of single-nucleotide polymorphism differences between mother and fetus [32]. Additionally, unlike liquid biopsy approaches using ultra-high-depth sequencing, the use of relatively low depth meaningfully decreases the cost of testing and permits the use of reasonable blood volumes, both of which will ultimately be required for population-level screening [12]. Finally, approaches focused on mutation detection alone can miss certain types of tumor-derived signals (e.g., genome-wide CNVs and epigenetic modifications), which are by definition most scarce in early-stage (i.e., non-metastatic) disease and pre-cancerous lesions, the detection of which is the goal of cancer screening programs.

While we have not yet directly determined the exact contributions to classifier performance from tumor-versus non-tumor-derived sources, several lines of evidence suggest that both may be present. First, while the observed relationship between AUC and inferred TF (Figure 3D) indicates that at least some of the classification power is likely attributable to the ability of the model to identify samples with abundant ctDNA, the ability to correctly classify samples with lower TF and/or early-stage disease suggests that ctDNA alone cannot fully account for classification performance. Second, a CRC classification model based solely on IchorCNA-estimated TF (inferred from CNV calls) performs relatively poorly, with an AUC lower than all tested CV results for the ML method, suggesting that non-CNV sources may contribute to our ML-based classifier. Future research will focus on better understanding the underlying biology of the classifier, as well as assessing potential improvements in model performance from the addition of other analytes and ML method development, including confounder mitigation.

In the presence of inadequately controlled confounders, ML methods are prone to learn irrelevant associations; this poses a critical challenge for the use of ML for biomarker discovery [33–36]. Certain confounders can be mitigated “up front” through experimental design (e.g., demographic biases and institution bias) or operational quality control (e.g., identification of known parameter drift). This can help minimize the dependence between class label and any potential noise-inducing variable but incurs an additional cost in time and/or operational expense. However, perfect control of confounders at the design stage is not realistic: Some variables may be intrinsically confounding in the population of interest (e.g., cancer incidence increases with age), and there are modes of variation which may exist but which may not be known a priori and therefore mitigated post hoc (e.g., batch-to-batch variability in sequencing).

A key contribution of this work is the presentation and analysis of cross-validation techniques specifically tailored to go beyond traditional k-fold validation to measure and mitigate a number of pervasive confounding effects in biomarker discovery: k-batch and ordered k-batch for different scales of process variability in time, respectively, and balanced k-batch for institution-specific biases. We found that standard k-fold CV can have higher performance than confounder-controlled CV methods, consistent with the historical difficulty in reproducing discovery studies. We believe that explicit stratification for technical and biological confounders may be used as standard practice to better evaluate the generalizability of early discovery results.

The current study has a number of potential limitations. First, because samples were obtained retrospectively, breaks in the chain of custody may have led to sample and labelling errors, which would impede the ability of an ML method to adequately learn. Additionally, the presence of CNVs in a small number of control samples (Supplemental Figure 2) has been previously observed in other cohorts and may be due to malignant or benign causes [5,14,37]; further follow up was not possible in this cohort. Another limitation of our study is that TF was estimated using copy-number inference from moderate-coverage whole-genome sequencing, which has a limit of detection of 3% for TF [28]; by contrast, targeted mutation detection would allow more sensitive characterization of TF. Furthermore, the presented cross-validation procedures control for individual confounders, but not for multiple simultaneous confounders; generalization of these procedures to multi-confounder control is an area for future work. Prospective studies are underway to validate classifier performance and verify generalization predictions from confounder-controlled CV.

Although this study focused on CRC, this study approach is directly applicable to other cancers and indeed to other pathological and physiological conditions. Our approach extracts signals from certain biological states and can apply them to better understand others; however, full development and validation of classifiers to address different clinical and non-clinical applications will require additional samples in those specific populations. Unlike targeted mutation approaches which require identification of disease-specific targets, this whole genome approach allows for the unbiased discovery of signals which are not disease-specific and could even be extended to the assessment and monitoring of non-disease states. Additionally, this approach should be able to detect unique epigenetic patterns for other diseases, thereby providing specificity by differentiating CRC from other cancers [38]. Efforts are currently underway to evaluate these hypotheses.

In summary, this study presents a novel representation of cfDNA and an analysis framework that demonstrates promising initial results for the detection of early-stage CRC based on a minimally invasive blood test. Prospective validation of this approach is currently underway, as is the incorporation of other cell-free, blood-based analytes (e.g., proteins) that may contribute orthogonal signals to further improve classifier performance.

## Supporting information

## Acknowledgements

The authors gratefully acknowledge Dr. Samuel So from the Asian Liver Center at Stanford University, Dr. Randall Brand from the University of Pittsburgh Medical Center, Dr. Andrew Godwin and the Biospecimen Repository Core Facility staff funded in part by the National Cancer Institute Cancer Center Support Grant (P30 CA168524), National Health Services Research Scotland, Tayside Biorepository, Geneticist Inc., iSpecimen Inc., and Indivumed for support of this research by providing de-identified plasma samples.

## Supplemental Information

Supplemental methods, figures and table are provided.

## Abbreviations

AUC: area under the curve
cfDNA: cell-free DNA
CHIP: Clonal hematopoiesis of indeterminate potential
CNV: copy number variant
CRC: colorectal cancer
ctDNA: circulating tumor DNA
CV: cross-validation
DNA: deoxyribonucleic acid
IU: intended use
ML: machine learning
RNA: ribonucleic acid
SVM: support vector machine
TF: tumor fraction

