## Supplementary material for "Machine learning enables detection of early-stage colorectal cancer by whole-genome sequencing of plasma cell-free DNA"

#### Supplemental Methods

##### Processing and sequencing

Cell-free DNA was extracted from 250 µl plasma (spiked with unique synthetic double-stranded DNA fragments for sample tracking) using the MagMAX cfDNA Isolation Kit (Applied Biosystems) per manufacturer instructions. Paired-end sequencing libraries were prepared using the NEBNext Ultra II DNA Library Prep Kit (New England Biolabs) with custom short Y-adapters containing unique molecular identifiers (UMIs), no "U Excision" step, and custom unique dual indexing P5 and P7 primers. Libraries were pooled in batches of 96 and sequenced on an Illumina NovaSeq 6000 Sequencing System across multiple S2 or S4 flowcells at 2x51 or 2x151 base pairs (bp), respectively, to a minimum of 400 million reads (median=636 million reads). After read trimming and UMI removal, effective read length for genome mapping of all samples was paired-end 45 bp.

##### Bioinformatics analysis

Reads were de-multiplexed and aligned to the human genome (GRCh38 with decoys, alt contigs, and HLA contigs) using BWA-MEM 0.7.15 (Li, 2013). Duplicate fragments were removed using fragment endpoints together with UMIs.

For all samples, sequencing data was checked for quality and rejected from downstream analysis if ≥ 1 of the following conditions were met: AT dropout > 10 (computed by Picard v2.10.5), GC dropout > 2 (computed by Picard v2.10.5), or deduplicated read pairs < 400 million. Additionally, we rejected samples in which the relative counts in sex chromosomes were inconsistent with the annotated gender. Finally, samples that were suspected of being contaminated (i.e., based on improbable observed allele fractions, spike-in cross contamination > 1%, low counts of spike-in reads, inter-sample contamination, or batches with a contaminated no template control) were manually verified prior to inclusion in the data set. Tumor fraction (TF) was estimated using IchorCNA (Adalsteinsson, 2017) using the parameters suggested for low tumor content samples (--normal "c(0.95, 0.99, 0.995, 0.999)" --ploidy "c(2)" --maxCN 3 --estimateScPrevalence FALSE --scStates "c()" --chrs "c(1:22)" --chrTrain "c(1:22)") and including a panel of normals comprised of healthy process control samples (disjoint from the healthy patient samples).

A cfDNA feature vector, or “profile”, was constructed for each sample by counting the number of fragments aligned to each annotated protein-coding gene in the genome. This type of data representation primarily captures two types of signals: (1) somatic CNVs (gene regions provide a sampling of the genome, enabling the capture of any consistent large-scale amplifications or deletions); and (2) epigenetic changes in the immune system at the level of nucleosome positioning as inferred by patterns in read pile-ups (Ulz, 2016), as nucleosome protection is believed to be the main mechanism for DNA preservation in plasma. Annotations for protein-coding genes (including both introns and exons but not the promoter region) were obtained from the Comprehensive Human Expressed SequenceS (CHESS) project (v1.0) (Pertea, 2018). Fragments were counted to produce a vector of “gene features” if the paired-end reads comprised a properly mapped pair, with at least one of the reads overlapping an annotated gene by one or more bases and the first read having a mapping quality of 60. This produced a vector with a dimensionality of 24,152 (covering 1,352 Mb) per sample. Each value in the vector corresponds to the number of reads that overlap a specific gene.

Preprocessing

We preprocessed the feature vectors via the following transformations: remove sex chromosomes, remove poor-quality features, and normalize the remaining features to account for length, read depth, and GC content. Poor-quality features were defined as those with mean mappability < 0.75 using a read length of 44bp (Derrien, 2012), GC percentage < 30% or > 70%, or reference-genome N content > 10%. For per-sample depth normalization, we used the trimmed mean (i.e., the mean after removing the bottom and top 10% of bins) as the scaling factor. We accounted for GC bias using Loess correction (Cleveland, 1979). Following these transformations, the resulting vector of gene features had a dimensionality of 17,582 features, covering 1,172 Mb.

Classification

The evaluated ML methods consisted of a series of transformations, in some cases including dimensionality reduction, followed by a supervised classification algorithm. A number of models were trained with k-fold CV; subsequently, the best performing model was evaluated with additional cross-validation procedures.

First, outliers (defined as feature values of a given sample that were above the 99th percentile of that feature across all training samples) were imputed to the 99th percentile value. Each feature was subsequently standardized across all training samples by subtracting the mean and dividing by the standard deviation. The same outlier replacement, using the means and standard deviations of the training set, were used to standardize the test set. If a dimensionality reduction transformation method was selected, it was trained on the training set and applied to all samples in both the training and the test sets. The dimensionality reduction transformations used in this study were truncated singular value decomposition (SVD) and principal component analysis (PCA).

Two possible classification algorithms were trained on the transformed input: logistic regression and support vector machine (SVM). Multiple hyperparameters were considered for each method using a random search of 100 iterations per fold with a validation set composed of 20% of the training data. Logistic regression had two hyperparameters: the inverse of regularization strength, and the choice of either L1 or L2 penalty. SVM had three hyperparameters: the inverse of regularization strength, tolerance for stopping criterion, and bandwidth of the radial basis function kernel.

Cross-validation

The purpose of a cross-validation (CV) procedure in assessing a classification model is to obtain an approximation or measurement of a model's performance on new, previously unseen data that were not used to construct the model. The goal is to provide an approximation by repeatedly training a model on a distinct subset of the data and testing on a held-out subset of data, unseen by the model during training. K-fold cross-validation procedure requires dividing the entire dataset into k groups. For each of the k groups (or folds), a model is trained with the other k-1 folds, and the held-out fold is used as the test set. Stratified k-fold cross-validation stratifies the samples by class before dividing into folds so that the approximate proportion of samples is roughly equivalent across folds.

##### Confounder analysis

In the presence of confounders, k-fold cross-validation may be susceptible to overestimating predictive performance because a statistical model may learn information about confounding covariates rather than the covariate of interest. That is, even though the original classification task is to predict the sample label (e.g., cancer versus non-cancer) from genomic feature data, the classifier may learn an association of confounders from the data. In such a case, classification results and estimated model performance may be misleading because the classifier learns undesirable associations between class label and the confounding factors within the training set and incorrectly applies these associations to the test set. To quantify the effect of each known confounder for which our dataset contains metadata, we evaluated the performance of a classifier trained to predict the sample label solely based on the confounding factor. Although it is unlikely any of our features are perfectly encoding confounding factors, these label confounding experiments represent the confounders' maximum contribution to performance. For these label-confounding experiments, each confounding factor was encoded as a categorical one-hot feature vector and used to train a model using k-fold cross-validation.

##### Controlling for confounding

Beyond simply quantifying the impact of each label confounder on model performance, there are ways to use cross validation to control for specific confounders. Confounders can be divided into at least two groups: those that are present in the sample, and those that are due to analytical processes. Controlling for known confounders can occur in one of two ways: 1) stratify the test sets so they contain only unseen elements of the confounding factor or, 2) control the distribution of the class labels for the confounding factor in the training set. The method selected depends on the data available.

Age is a confounder inherent to our study population because cancer incidence is known to increase with age (Cronin et al, 2018). This is evident in preliminary analyses of age distributions by class. Therefore, binned-age CV assesses the impact of patient age on model performance by training and testing on samples from patients in different age ranges. To create binned ages, samples from patients with known ages were grouped into 6 bins within the IU age range (i.e., [50-59], [60-64], [65-69], [70-74], [75-79], [80-84]). During model training, each age bin corresponds to one fold, and the model is trained using 5 of the 6 folds; the remaining fold is used for testing.

Analytical confounders are divided into short-term, long-term, and institution/site effects. Short-term effects are those that co-occur within a batch of samples and can be directly attributed to the batch on which the samples were processed. To control for these effects, k-batch CV stratifies samples by batches; that is, samples in the test set will not come from a batch that was also used during training.

Another confounder, particularly in studies of retrospective samples collected from many sources, is institutional or site-to-site variability in sample collection and pre-analytic processing protocols. Variation in sample processing protocols can be approximated by grouping samples by source institution. When grouping samples in this manner, the distribution of class labels can be extremely imbalanced due to different types of patient samples being obtained from different institutions. This makes cross-validation by source institutions incomparable to models trained with k-fold. Therefore, to enable a direct comparison of model performance based on holding out samples from different source institutions, training samples are downsampled so that samples from each institution have the same distribution of class labels. Balanced k-batch CV applies institutional downsampling to folds stratified by batches.

Finally, k-batch cross-validation works well for controlling biases within a batch, but analytical processes can drift over extended period of time despite still passing QC. The sources are vast but can include changes in laboratory equipment, sample processing procedures, and operator personnel. Such variability can also be a confounder. In this study, cancer and non-cancer samples were processed over approximately one year, so there was the potential for process drift. To assess the potential effects of process drift on model performance, ordered k-batch sorts batches by processing date and divided into folds similar to a time series split of time-dependent data.

For CRC, samples without a known age or whose age was outside the IU age range were included in all training folds but no test folds; all other samples were partitioned to appear exactly once in testing in each cross-validation procedure. Model performance was therefore evaluated only on samples from the IU age range, consistent with commercially available CRC screening tests [39].

### Supplemental Results

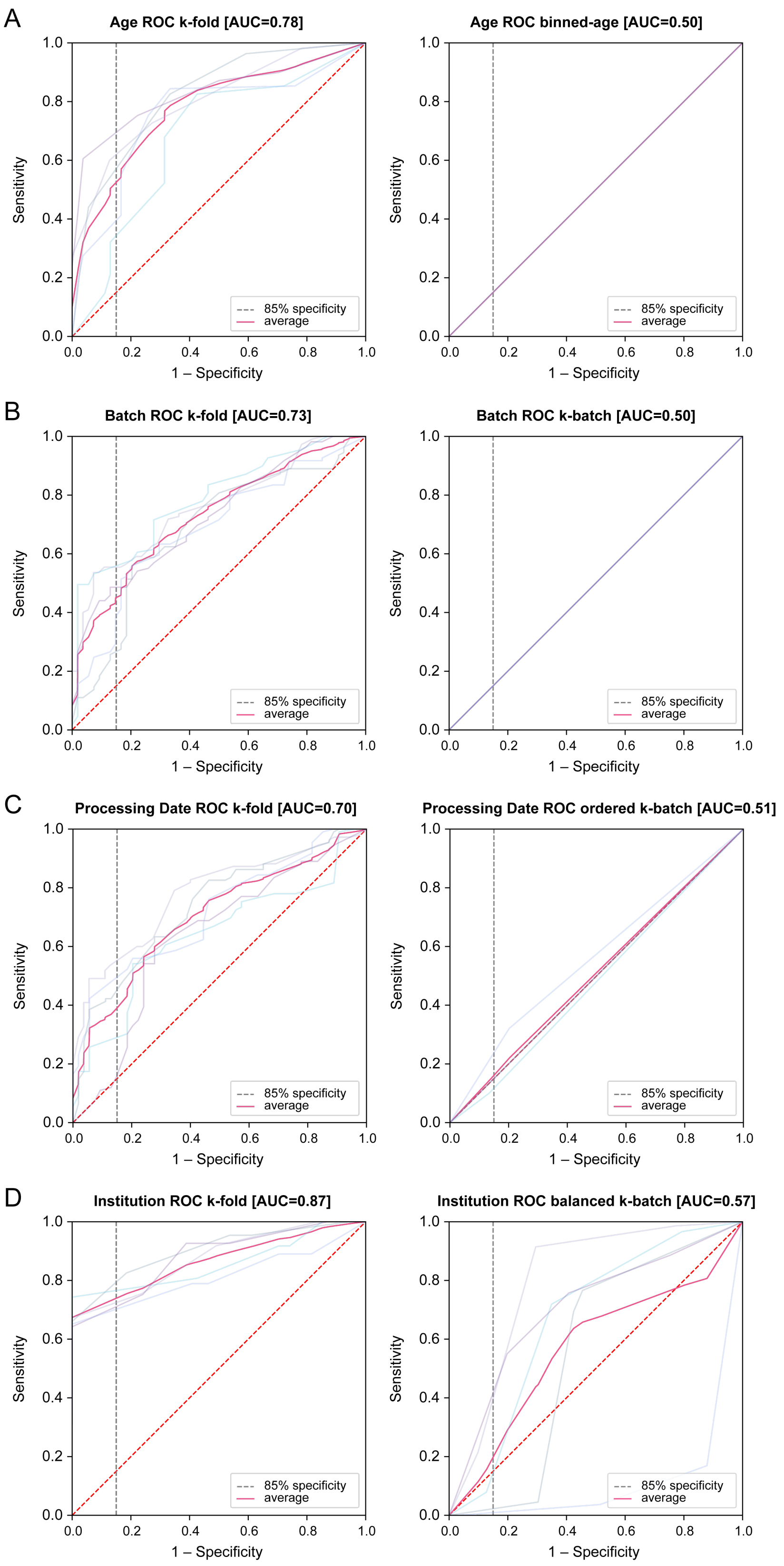

**Supplemental Figure 1:** Performance of confounding variable in k-fold and target CV procedure (ROC curves) for:

- **A.** Age and binned-age CV
- **B.** Batch and k-batch CV
- **C.** Processing date and ordered k-batch CV
- **D.** Institution and balanced k-batch CV

Note that stratification CVs (i.e., binned-age, k-batch, and ordered k-batch) perform at chance (mean AUC of 0.50) when the distribution of labels in the test covariate is even and would result in an ROC that is just a diagonal line. The variance seen in some of the target procedures is due to class imbalance in the test data.

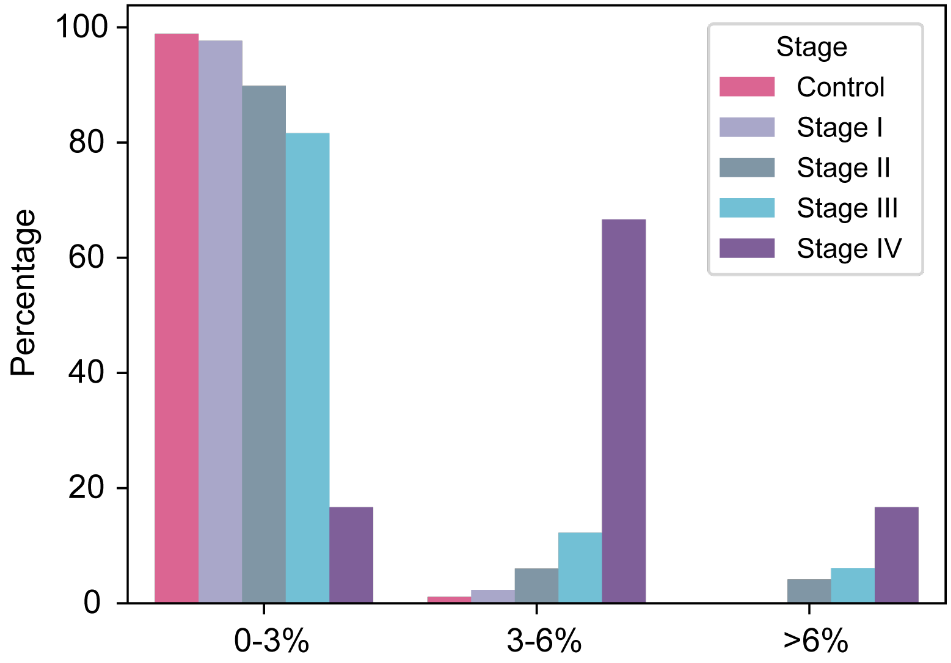


**Supplementary Figure 2:** Percentage of CRC samples by stage and non-cancer controls in each range of estimated TF (which computed from observed CNV using IchorCNA).

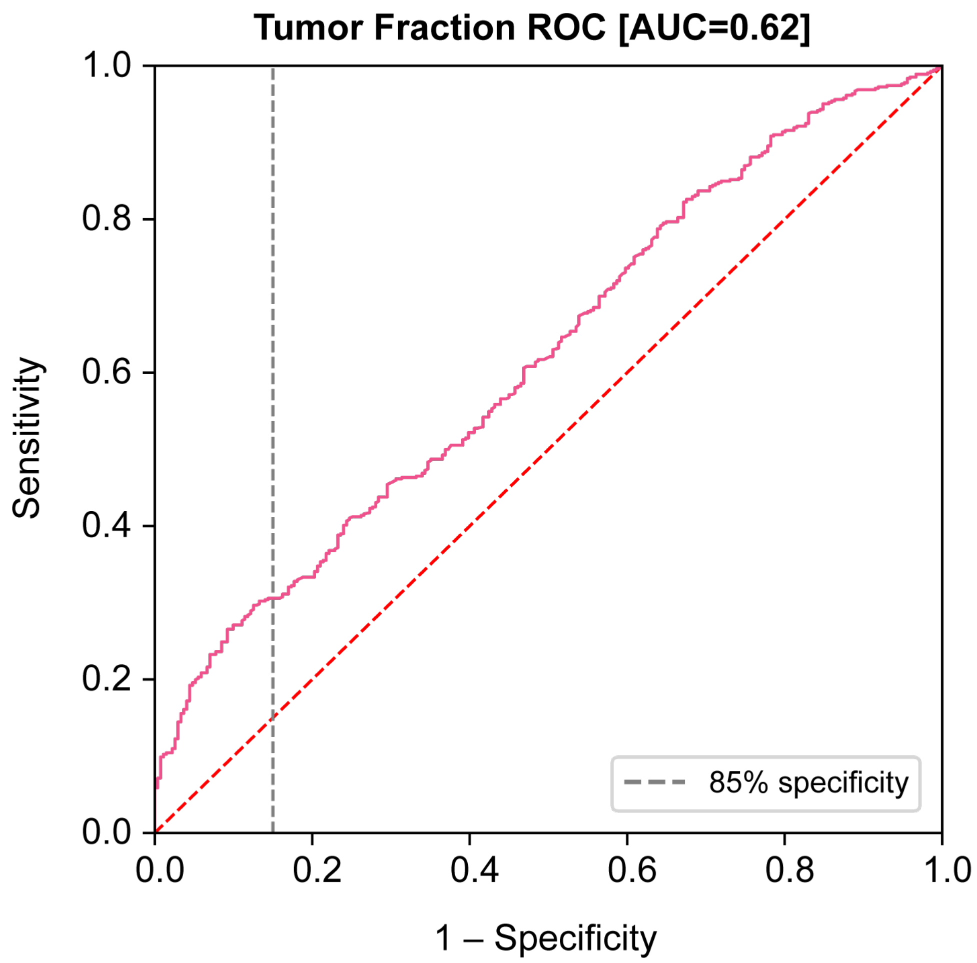


**Supplemental Figure 3:** IchorCNA-estimated TF alone to predict cancer achieved an AUC of 0.62 in the IU age range, meaningfully lower than results from the ML model under any analyzed CV procedure.

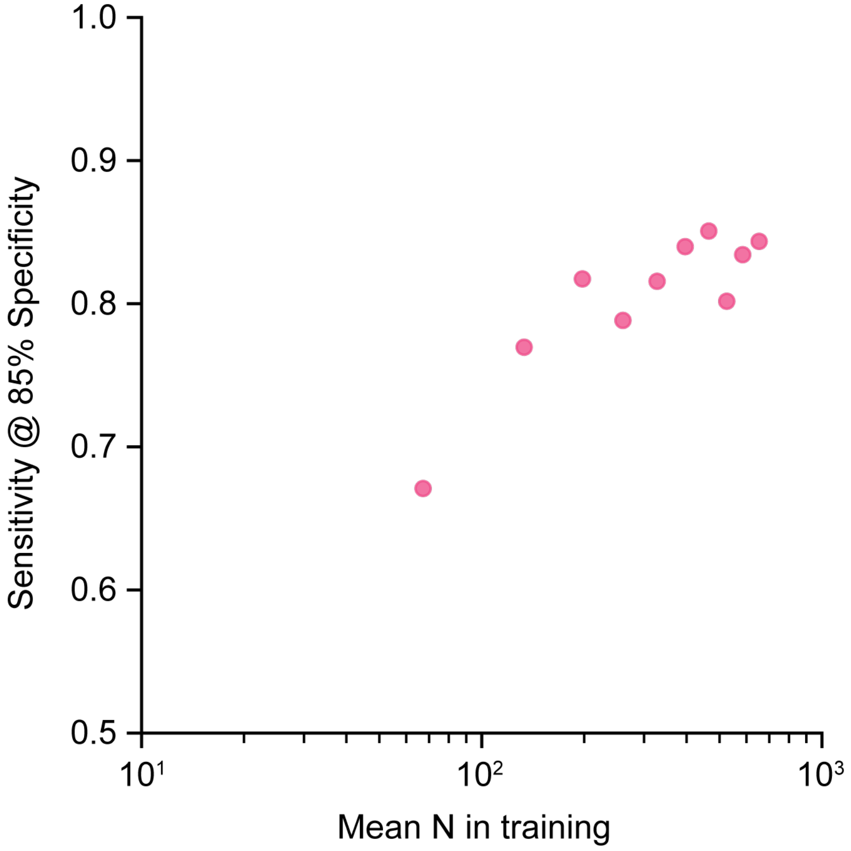


**Supplemental Figure 4:** Non-linear relationship between the total number of samples used for training and AUC for CRC detection. The method selected by CRC analyses was trained again with k-fold except the number of training samples per fold were downsampled. The lower numbers are comparable to those available for balanced k-batch and were used to investigate decreased classifier performance due to smaller sample sizes in training.

| | Number of folds | Mean N in Training | SD of N in Training | Mean N in Test | SD of N in Test |
|---|---|---|---|---|---|
| **k-fold** | 5 | 653.6 | 0.89 | 163.4 | 0.89 |
| **Binned-age** | 6 | 707.8 | 26.4 | 109.2 | 26.4 |
| **k-batch** | 5 | 653.6 | 35.0 | 163.4 | 35.0 |
| **Ordered k-batch** | 5 | 653.6 | 80.8 | 163.4 | 80.8 |
| **Balanced k-batch** | 5 | 263.6 | 39.0 | 163.4 | 35.0 |

**Supplemental Table 1:** Mean and standard deviation of the number of samples (both in and out of the IU age range) per fold for each CV procedure. SD = standard deviation
